# Imaging of neural oscillations with embedded inferential and group prevalence statistics

**DOI:** 10.1101/164731

**Authors:** Peter W. Donhauser, Esther Florin, Sylvain Baillet

## Abstract

Magnetoencephalography and electroencephalography (MEG, EEG) are essential techniques for studying distributed signal dynamics in the human brain. In particular, the functional role of neural oscillations remains to be clarified. Imaging methods need to identify distinct brain regions that concurrently generate oscillatory activity, with adequate separation in space and time. Yet, spatial smearing and inhomogeneous signal-to-noise are challenging factors to source reconstruction from external sensor data. The detection of weak sources in the presence of stronger regional activity nearby is a typical complication of MEG/EEG source imaging. We propose a novel, hypothesis-driven source reconstruction approach to address these methodological challenges^1^. The imaging with embedded statistics (iES) method is a subspace scanning technique that constrains the mapping problem to the actual experimental design. A major benefit is that, regardless of signal strength, the contributions from all oscillatory sources, which activity is consistent with the tested hypothesis, are equalized in the statistical maps produced. We present extensive evaluations of iES on group MEG data, for mapping 1) induced oscillations using experimental contrasts, 2) ongoing narrow-band oscillations in the resting-state, 3) co-modulation of brain-wide oscillatory power with a seed region, and 4) co-modulation of oscillatory power with peripheral signals (pupil dilation). Along the way, we demonstrate several advantages of iES over standard source imaging approaches. These include the detection of oscillatory coupling without rejection of zero-phase coupling, and detection of ongoing oscillations in deeper brain regions, where signal-to-noise conditions are unfavorable. We also show that iES provides a separate evaluation of oscillatory synchronization and desynchronization in experimental contrasts, which has important statistical advantages. The flexibility of iES allows it to be adjusted to many experimental questions in systems neuroscience.

**Author summary:** The oscillatory activity of the brain produces a repertoire of signal dynamics that is rich and complex. Noninvasive recording techniques such as scalp magnetoencephalography and electroencephalography (MEG, EEG) are key methods to advance our comprehension of the role played by neural oscillations in brain functions and dysfunctions. Yet, there are methodological challenges in mapping these elusive components of brain activity that have remained unresolved. We introduce a new mapping technique, called imaging with embedded statistics (iES), which alleviates these difficulties. With iES, signal detection is constrained explicitly to the operational hypotheses of the study design. We show, in a variety of experimental contexts, how iES emphasizes the oscillatory components of brain activity, if any, that match the experimental hypotheses, even in deeper brain regions where signal strength is expected to be weak in MEG. Overall, the proposed method is a new imaging tool to respond to a wide range of neuroscience questions concerning the scaffolding of brain dynamics via anatomically-distributed neural oscillations.

## Introduction

The role of neural oscillations in population codes of brain functions, and the possible mechanisms of inter-regional communication between brain regions are not entirely understood. Source imaging techniques with magnetoencephalography (MEG) or electroencephalography (EEG) are time-resolved, non-invasive tools used to test a great diversity of neurophysiological hypotheses (1). In principle, MEG/EEG imaging can map multiple regional sources of oscillatory activity from external sensor data. However, spatial smearing and heterogeneous signal strength across brain locations limits the performance of current source imaging methods. Consequently, if nearby brain regions express an effect of interest, the area of stronger magnitude will mask the detection of weaker sources, as illustrated in Fig 1. The detection of multiple oscillatory sources therefore remains challenging to MEG/EEG imaging. This limits the insight about distributed brain dynamics that can be gained from the technique.

**Fig 1.**
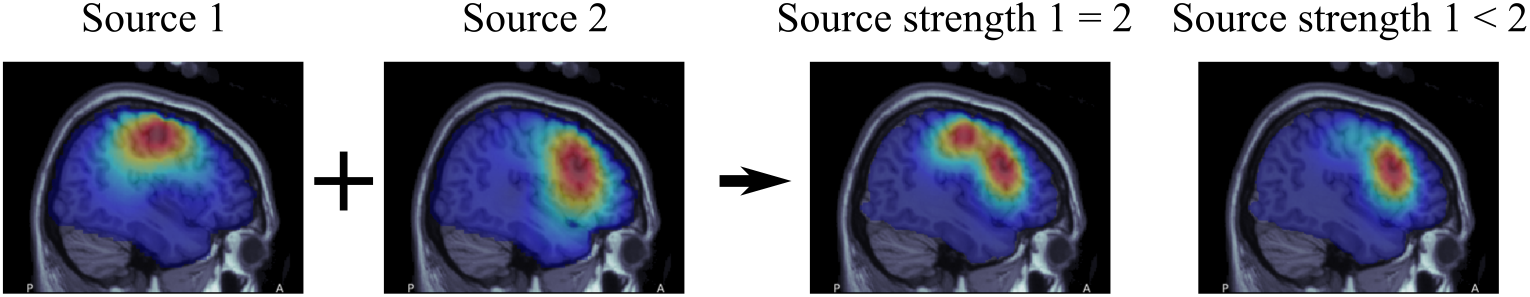
Illustration of the field-spread effect on the detection of weak MEG sources. Point-like sources are typically recovered using beamforming or minimum-norm estimation (MNE) imaging at the expense of exaggerated spatial smearing in the source space. The Source 1 and Source 2 maps are examples of such effect when active separately. When both sources are active simultaneously, their relative strengths impact the ability of source imaging to spatially resolve between the two active regions. When both source magnitudes are similar (Source strength 1 = 2), the map can display their respective contributions. When one source is weaker than the other (Source strength 1 *<* 2), its presence in the resulting source map may be masked by that of the strongest source (here Source 2).

MEG/EEG localization of oscillatory generators typically relies on a procedure that is non optimal in terms of signal detection. Source time-series are first reconstructed using imaging or beamforming approaches (2). Second, inferential statistics based on the experimental hypothesis are tested at each voxel of the source space – e.g., using the ratio of oscillatory power between two experimental conditions. Significant and spatially-distinct regional clusters are then interpreted as distinct sources of oscillations. This approach hinders the detection of weaker or deeper sources in the presence of stronger regional activity. We refer to this methodology as the *standard approach*.

We introduce a novel methodology to alleviate this problem. The technique performs imaging with embedded inferential and group prevalence statistics (iES) altogether. With iES, the experimental hypothesis is not deferred to the stage of statistical inference on the estimated source values. Rather, it explicitly constrains the solution to the hypothesis tested. In essence, iES reduces the (spatial) dimensions of the data, to detect and equalize the contribution of source components that are consistent with the tested hypothesis. The iES methodological apparatus is based on generalized eigen decompositions (see e.g. 3), nonparametric statistics (4; 5) and subspace scanning (MUSIC, see 6). We explain here that iES 1) has key statistical advantages, yielding improved detection sensitivity, 2) can be used in conjunction with the standard approach, for complementary estimation of source strengths, 3) improves the detection of functionally connected regions, and that 4) iES can implement a wide range of experimental hypotheses.

## Results and Discussion

### Overview of the approach

We describe the basic principles of iES and illustrate the steps involved using a MEG data example. The method per se is detailed in Materials and Methods.

#### Basic principles

We propose to transcribe the experimental hypothesis into a quality function *f* (s) over a signal **s**. *f* (s) is defined such that it returns larger values if s is consistent with the hypothesis. In Fig 2a we show the quality functions featured as examples in this article, with multiple possible variations, as discussed below. For example, in the case where stimulus-induced responses in the gamma band (50-85 Hz) are expected, *f* (s) would be designed to return the ratio of gamma power between time segments when the stimulus is presented vs. when the stimulus is absent. Note that the tested design is a directed one: in the latter example, testing for gamma power increases vs. decreases are two different hypotheses that are evaluated separately. Fig 2a) shows several use cases and signals s that are either consistent or inconsistent with the hypothesis quantified by *f* (s) values.

**Fig 2.**
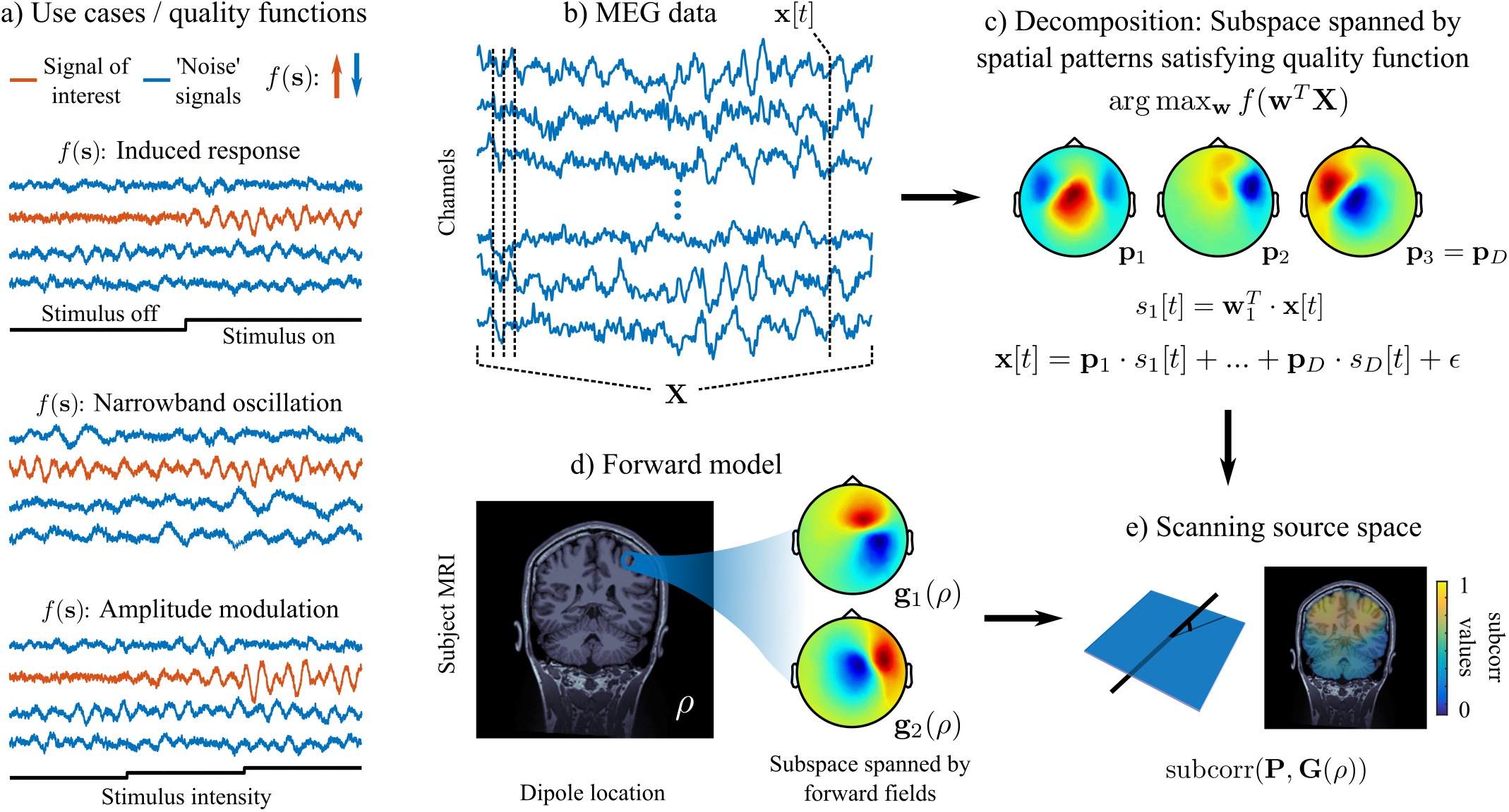
Basic principles of iES. ***a) Examples of designs:*** The experimental design (shown as a black trace) determines the quality function *f* (**s**), so that this latter takes high values for signals consistent with the hypothesis (in orange; the signals that do not correspond to the tested hypothesis are shown in blue). *b) MEG data:* the multichannel MEG recordings are captured in the matrix **X** = {**x**[*t* = 1], *…,* **x**[**t** = **T**]}. *c) Computing the signal subspace:* spatial patterns **P** = {**p**_1_, *…,* **p**_*D*_} are extracted from the MEG data by optimizing the quality function with respect to spatial filters **W** = {**w**_1_, *…,* **w**_*D*_}. Whereas **W** is used to extract the signals of interest from the multichannel MEG data, **P** are the forward fields of these signals as they contribute to the measured MEG data. *d) Computing the forward model:* shown are the MEG spatial patterns **G**(*ρ*) generated by two tangential dipoles at location *ρ* in a single subject. *e) Subspace correlation as a scanning metric:* The spatial patterns from c) and d) span a subspace of the MEG sensor space. A grid of source locations is scanned with a subspace correlation metric (6), quantifying the smallest possible angle between the data and source subspaces. This yields a distributed map of scores, which highlights possible source locations consistent with the hypothesis.

Let x[*t*] denote the MEG/EEG time-series recorded from an array of channels (Fig 2b) and **X** the data matrix (channels *×* time samples *t*). The quality function *f* is used in an optimization problem to identify spatial filters **w**_*i*_ and spatially-filtered signals *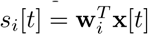* in the data such that the quality function is maximized as

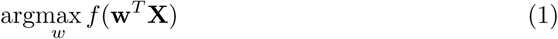

and spatial patterns **p**_*i*_, *i* = 1, *…, D* describing the corresponding topographies that contribute to the recorded time-series as

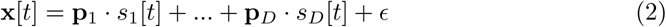

(Fig 2c). The combination of spatial filters, patterns and corresponding signals is comparable to the notion of ‘components’ in independent component analysis (ICA, 7), which yields mixing (spatial patterns **p**_*i*_) and unmixing (spatial filters **w**_*i*_) matrices as well as ICA time-series (spatially filtered signals *s*_*i*_[*t*]). The spatial patterns of signals that conform to the tested hypothesis (i.e. have high quality function values *f* (**s**)) represent a subspace of the MEG/EEG channel space (the signal subspace, see 6).

The MEG/EEG forward model predicts spatial patterns **g**(*ρ*) produced at the channel array by an elementary current source at location *ρ* (Fig 2d). Therefore the forward fields of current sources with different orientations also form a subspace of the MEG/EEG channel space. The iES method proceeds by scanning each elementary brain location of the source space. The source space can be a uniform 3-D grid of the brain volume, or restricted to the cortical surface (Fig 2e). At each tested brain location, the correspondence between the forward fields from this location and the data spatial patterns identified by the quality function is evaluated using the measure of subspace correlation (*subcorr*). This latter quantifies the smallest principle angle between two subspaces (6). Intuitively, the data and the physical forward models are compared at each brain location, with respect to the experimental hypothesis. The procedure generates a map of possible sources, which activity accounts for the experimental effect of interest.

Since the effect strength is entirely captured in the quality function *f* (s), it does not directly influence the *subcorr* values. Therefore, the contribution of the hypothesis-consistent sources are equalized in the resulting maps, which contributes to their detection, regardless of their respective strengths.

#### Extension to group-level analysis

So far, subspace scanning techniques have been mostly used to identify sources of event-related trial averages in single-subject MEG/EEG data (6; 8). We describe a principled approach to conduct group level analyses with the proposed method, using an example data set. The data was obtained from a variation of the visual attention experiment in (9), where a contracting circular grating was presented to participants in MEG. In each trial, after 3-5 seconds following stimulus onset, a change in the contraction speed occurs and participants had to indicate their perception of the change with a button press. To illustrate the methodology, iES was used to identify the regions where gamma-band (50-85Hz) power was stronger during visual stimulus presentation, with respect to baseline, prestimulus periods. Gamma oscillations generated in occipital visual regions are expected to be reliably enhanced by this paradigm (9). The quality function *f*_*induced*_ was defined as the ratio between the gamma power during the interval [1, 3] seconds post-stimulus onset, and the gamma power during the baseline interval [-2, 0] seconds (0s corresponds to stimulus onset).

Fig 3 shows the iES group analysis workflow in detail. First, the signal subspace estimation described above yields components from each participant that can be interpreted similarly to those of a principal component analysis (PCA) decomposition (Panel A). The subset of spatial patterns retained for source-space scanning corresponds to the iES components with highest quality function scores, exceeding a threshold *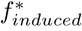*. This threshold is determined by a permutation procedure under the null hypothesis of exchangeability of baseline and stimulus data segments. A permutation histogram of *f*_*induced*_ values is obtained, and *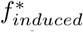* is set to the value that is higher than 1–*p*_*crit*_*/*2 values of the permutation distribution, with *p*_*crit*_ = .05 in the example presented. In our example, this procedure yields five spatial components to be included in the definition of the signal subspace (Fig 3a). The sensor spatial patterns suggest occipital signal origins of stronger gamma-band activity during stimulus presentation. The iES decomposition, akin to PCA, produces orthogonal signal components s_*i*_. Thus the corresponding spatial patterns do not necessarily represent anatomically distinct sources: the spatial localization of corresponding sources is subsequently obtained via scanning of the source space.

**Fig 3.**
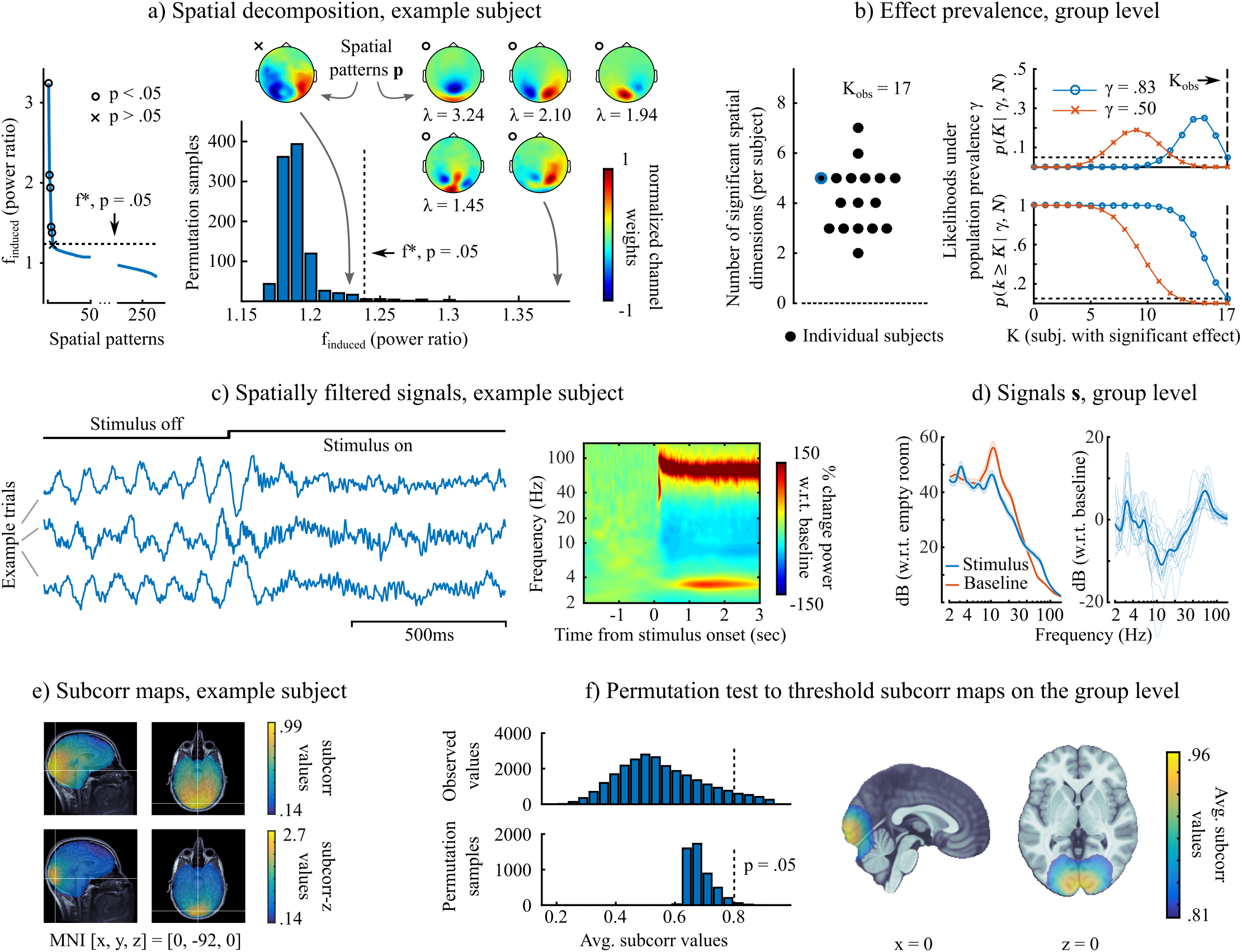
iES group analysis: mapping induced gamma oscillations during visual stimulation. *a) Subspace computation, example subject:* (left) values of the quality function *f*_*induced*_ for all the spatial patterns in the MEG data, ranked in decreasing order. The components with the 5 largest values of the quality function were deemed consistent with the tested hypothesis (highlighted with black dots - left, and their sensor topographies shown to the right). This was determined via permutation tests, which yielded *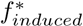*, a threshold indicating the minimum value of the quality function for significance (*p <* 0.05). Note that the number of significant components may vary per subject, as illustrated hereafter. *b) Effect prevalence, group level:* (left) number of significant spatial components for each subject (*K*_*obs*_ = 17 is the number of participants in this example). The subject illustrated in Panel A is shown in blue; (right) prevalence testing results (as detailed in Materials and Methods) showing the likelihood of the data under a population prevalence γ γ = .83 is the highest value that can be rejected at *p <* 0.05. *c) Spatially-filtered signals, example subject:* (left) three example trials: the increase in gamma oscillations after stimulus presentation can be readily appreciated visually in the spatially-filtered signals; (right) average time-frequency map across 220 trials: here too, the strong induction of gamma activity is clearly visible. *d) Spatially-filtered signals, group level:* (left) average power spectra of spatially-filtered signals in the two experimental conditions (baseline and visual stimulus) are scaled with respect to empty-room MEG recordings, shaded regions are standard errors; (right) the difference of the power spectra between the two conditions, with thin lines representing single-subject data. *e) Subspace correlation maps, example subject:* (top) Map of *subcorr* values in the 3-D source grid, indicating the location of brain regions generating stimulus-induced gamma activity, (bottom) Fisher-z transformed map. *f) Subspace correlation maps, group level:* (left) a permutation procedure to determine a statistical threshold to apply on the average *subcorr* scores. The figure shows the histograms of the permuted and observed *subcorr* values; (right) group-level average *subcorr* map, thresholded at *p <* 0.05. The effect confirms the single-subject data shown, and localizes to the occipital visual cortex.

At the group level, our approach acknowledges that the effect being tested may be absent in some participants. Concretely, their data may not contain a spatial pattern whose *f*_*induced*_ exceeds the critical value 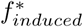. Rather than pretending otherwise and averaging across all participants, as is done in the standard approach, we put forward the concept of population prevalence γ to account effectively for the variability of the tested effect in the group (Fig 3b) (see also 10, for similar discussions). This notion enables to form a prediction on how many subjects in the sample are expected to show an effect. A prevalence null hypothesis, *H*_0_ : γ *≤* γ_0_, can be tested using a simple binomial distribution. The null hypothesis can be e.g., that the effect is absent from the population (γ_0_ = 0, global null hypothesis) or that it is present in less than half of the population (γ_0_.*≤* 5, majority null hypothesis). The null hypothesis is rejected if observing the number of subjects presenting the effect has a probability lower than a critical value (here *p*_*crit*_ = .05). In the present example of induced gamma oscillations, all subjects in the sample show the effect of interest. This means we can reject the majority null hypothesis (γ_0_ *≤* .5), and the highest γ_0_ that can be rejected at the given significance level is γ_0_ = .83, which can be interpreted as a lower-bound estimate on the population prevalence.

The spatial patterns **p**_*i*_ in Fig 3a all have corresponding spatially-filtered signals *s*_*i*_[*t*]. We show in Fig 3c and d examples of time series, time-frequency decompositions and power spectra of the signals corresponding to the largest effect (*s*_1_[*t*]). By construction (in terms of maximizing the *f*_*induced*_ quality function), these signals present the largest ratio of gamma power between stimulus and baseline periods. Strong, tonic gamma oscillations are clearly visible after stimulus onset, along with reductions in alpha/beta power (9). A 3.2-Hz oscillatory component is also found: it corresponds to the entrainment of lower-frequency neural components at the pattern-repetition frequency of the contracting circular grating.

We used the *subcorr* metric to produce source-level maps for each effect-prone subject, i.e. whose data features a non-null signal subspace. We show an example of an individual *subcorr* map in Fig 3e), which as expected, indicates a spatial peak in occipital visual regions. The Fisher-z transform arctanh(*subcorr*) can be applied to obtain a sharper map (referred to as *subcorr*-z). We then performed statistical inference at the group level, using group averaged *subcorr* maps in a permutation procedure (Fig 3f). A permutation distribution of the maximum-statistic is computed under the null hypothesis of exchangeability of signal subspace with a dimension-matched subspace drawn from the opposite end of the decomposition spectrum in Fig 3a. The null hypothesis thus reflects the assumption that the effects were not localized and spatially consistent across the tested cohort. This procedure yielded a statistically thresholded map of average *subcorr* values, highlighting the brain regions spatially consistent across the group, with an activity profile responding to the experimental question of interest. Here, the resulting map pointed to the visual cortex as the source of the gamma oscillations induced by the visual stimulus. This result was expected from published reports, and therefore further strengthens the validity of the proposed approach.

#### Distinct evaluation of positive and negative effects improves statistical power

The iES source maps highlight sources whose signals are consistent with a directed hypothesis across a group of subjects. When two experimental conditions are contrasted, this implies that two distinct source maps can be produced: for instance in the previous case example, one map corresponding to increased oscillatory power in one condition over the other; the other map corresponding to decreased oscillatory power. The benefit resulting from this is that mutual interference in the detection and statistical evaluation of the two sets of sources is avoided.

We demonstrate these methodological assets using the same experimental MEG data as above. We analyzed task-induced oscillations in the beta band (13-30 Hz), with the hypothesis that they were strongly suppressed during attention-demanding tasks in the occipital visual cortex (9; 11). We also wished to test whether other brain regions would reveal a selective increase in beta power during stimulus presentation. This contrast thus serves to illustrate how a strong power effect (decreased beta power) can challenge the detection of weaker opposite responses (increased beta power) with the standard approach, but not with iES.

Figure 4 shows results for the hypothesis of increased beta band power during stimulus presentation. The data from an example subject (Panel a) contained one spatial component consistent with that hypothesis. At the group level, only eight subjects out of 17 showed the effect of interest. Here, we shall emphasize the importance of the notion of effect prevalence, since the majority null hypothesis could not be rejected (Fig 4b). However, the prevalence null hypothesis can be rejected up to γ_0_ = 0.22, which indicates there is a subgroup of the population from which our subjects were drawn, which show the hypothesized effect. To better illustrate the significance of this notion, let us first assume the effect is not present in the population. With a probability of 0.95, one may still observe out of chance an effect in up to 3 out of the 17 subjects. The prevalence test therefore indicates that the observed data is unlikely under the assumption that prevalence is 22% or less (at a false positive rate of *p <* .05). Thus we pursued further the analysis of the subgroup of 8 participants (see Fig 4b), bearing in mind that the results may not generalize to the majority of the population. The validity of such a decision depends on whether the scientific question is pertinent to generic vs. restricted effects among participants. For instance, it can be particularly valuable for identifying effects that are more specific of a sub-type of participants in terms of behaviour or clinical condition.

**Fig 4.**
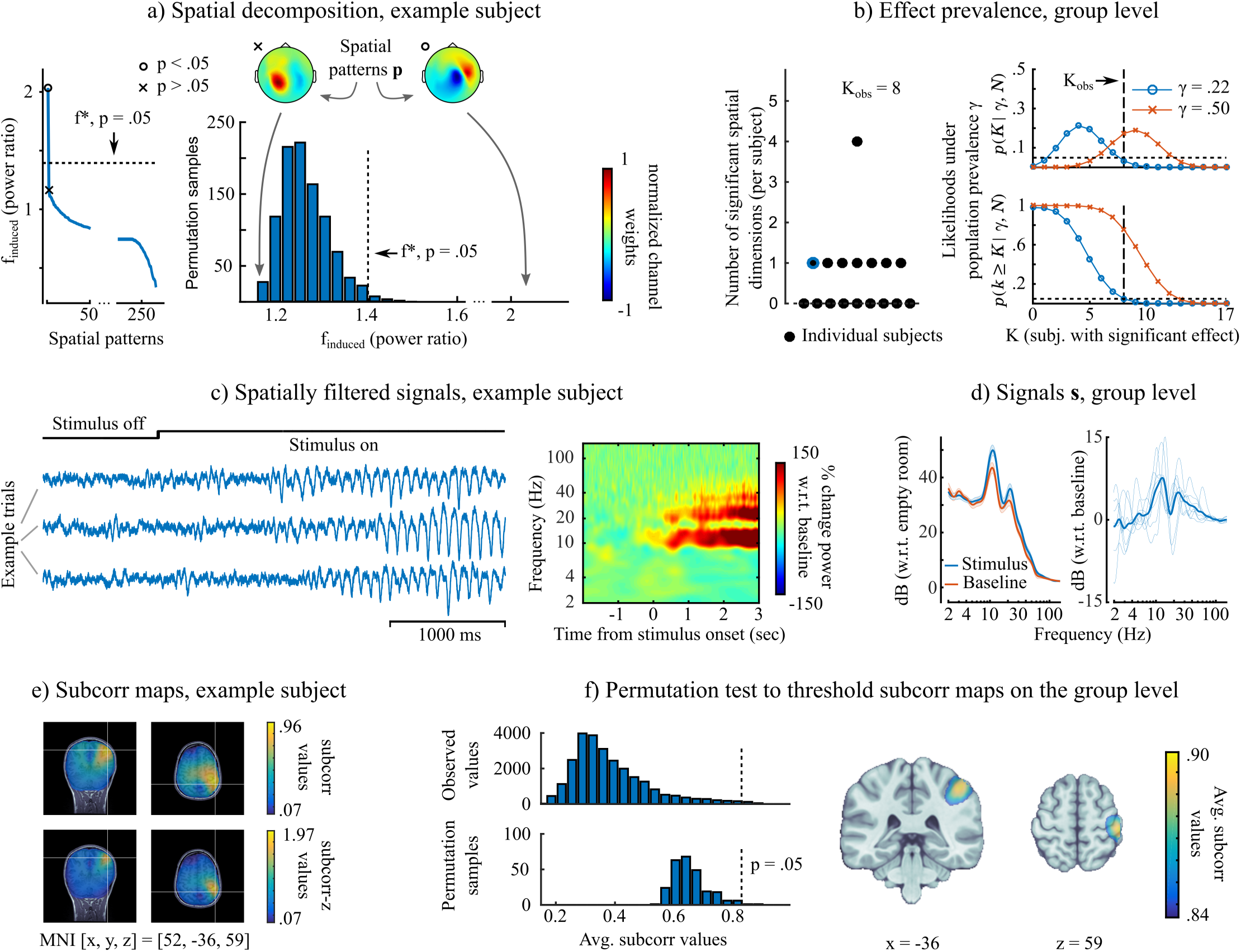
Effects in a subgroup of participants: mapping induced oscillations in the beta band (13-30 Hz) during visual stimulation. The data is that of Figure 3, and the present figure layout is identical. *a) Subspace computation, example subject:* in this participant, only one significant spatial dimension was retained for the signal subspace contributing to stronger power in the beta band. *b) Subspace computation, group level:* γ = 22 was the highest population prevalence that could be rejected at a *p* = 0.05, thus the majority null hypothesis could not be rejected. The analysis was pursued with the subgroup (n=8) of participants that showed the hypothesized effect. The purpose was to appreciate the spatial concordance across subjects and compare iES to standard source imaging approaches. *c) Spatially filtered signals, example subject:* induced power changes in the band of interest (beta, but also in alpha band) are clearly visible in 3 example trials. *d) Spatially filtered signals, group level:* induced power changes in the band of interest were found in the participant subgroup (n=8). *e) Subspace correlation maps, example subject:* the hypothesized effect localized to the right post-central/parietal cortex. *f) Subspace correlation maps, group level:* the effect localized to the right post-central gyrus. Note that this effect cannot be generalized to the majority of the population that the subjects were drawn from (see b) but only to a subset, which may present interesting capacity for identifying subtypes in participants.

Fig 4c-d shows typical signal traces in a subject from the subgroup presenting stronger beta and alpha oscillations building up during stimulus presentation. The sharp waveforms and the combined alpha/beta spectral pattern were typical of the somatosensory mu rhythm (12; 13). The effect was localized to right postcentral regions, as shown in the example subject and the group *subcorr* maps (Fig 4e-f). This result replicates previous observations of lateralized beta oscillations during an attention-demanding task (14).

To compare these findings with those from the standard approach, we obtained source maps of log-power ratios using minimum-norm imaging kernels. We used the MNE implementation of Brainstorm, with default parameters (15). The resulting maps were statistically thresholded following the same permutation procedure based on the maximum statistic. Note that with this procedure, distinct maps of positive and negative effects cannot be produced. For comparison purposes, we used the data from the subgroup (n=8) that showed the desired effect of higher beta power during stimulus presentation.

Fig 5a shows the complete beta band iES results (i.e. increases and decreases). In addition to the increased stimulus-induced beta power over right postcentral regions, we observed beta suppression in the visual cortex. The prevalence assessment revealed that this latter effect was observed in the entire group, and thus may generalize to the majority of the population. In the minimum-norm map (Panel b), the suppression of beta oscillations in the visual cortex was also readily observed, with similar spatial extension. However the increased, stimulus-induced beta oscillations over the right central regions were absent from the minimum-norm map produced from the 8 subjects presenting the effect in iES. The non-thresholded maps are shown in Supplementary Material, and confirm that a positive peak was indeed present in the minimum-norm maps, but was not deemed statistically significant. The reason for the observed discrepancy between methods can be understood from the permutation and data histograms (Fig 5, right column): By definition, the permutation histograms of the log-power ratios were mirror images for the evaluation of positive and negative effects respectively. This was the case because we drew exhaustive permutations from the data from the 8 subjects (28 = 256). Thus every unique permutation of labels had a corresponding opposite permutation. The consequence was that the variance and spread of the resulting distribution were determined by the strongest effect in magnitude – here the negative effect of beta suppression. The histogram of the observed data indicated that the right tail of the histogram indeed did not reach the statistical threshold. The iES allowed to test two directed hypotheses separately. Hence the permutation distributions were distinct and adapted to each respective hypothesis, revealing the positive effect in the iES statistical source map that were absent in the standard approach.

**Fig 5.**
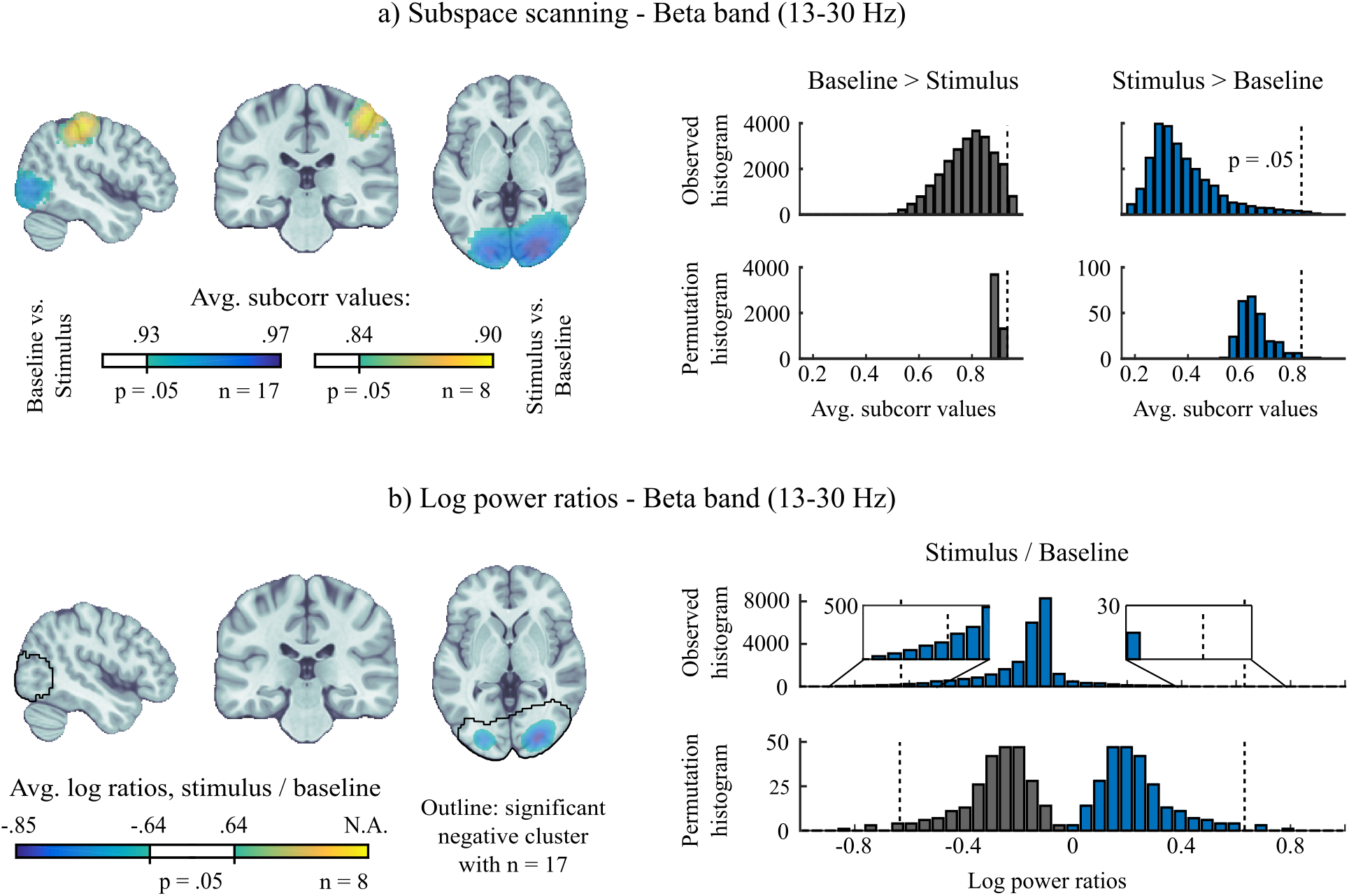
Comparing statistical performance of iES with minimum-norm imaging for mapping induced oscillations. *a) Subspace scanning results:* (left) significant average *subcorr* map (*p <* .05, see text for procedure). Note that results were obtained from the subgroup of participants that presented the hypothesized effect (n=8, see Fig 4). (right) histogram from observed data and permutation tests to derive a *subcorr* threshold corresponding to *p <* .05. *b) Minimum-norm imaging results:* average maps of log-transformed power ratios (stimulus/baseline, *p <* .05). Note that the distinction between positive and negative effects is not possible. The results were derived from the same subgroup (n=8) to allow comparison with a), the results obtained with the full group (n=17) are shown as an outline. Contrary to iES, no increase in beta power could be detected over the right post-central gyrus region, with the same subgroup of subjects. Unthresholded maps are shown in the supplementary material. (right) histograms of observed data and permutation tests to determine significance of minimum-norm maps at *p <* .05. Note how the strong negative effects inflated the permutation distribution and prevented the detection of the smaller positive effects. As shown using iES, positive and negative effects could be evaluated separately and specifically.

#### Supplementary insight gained compared to standard approaches

We detail in Methods that iES requires the computation of cross-spectral or covariance matrices, and their decomposition in the generalized eigenvalue framework. This means that in addition to the *subcorr* statistical maps produced, a corresponding map of the standard approach can be obtained by applying a minimum-norm imaging kernel to those matrices, which allows plotting the value of the quality function *f* at each location of the source grid. Fig 5 shows an example of this approach to obtain a map of log-power ratios. We emphasize that the combined use of *subcorr* and minimum-norm source maps enabled by the proposed method provides complementary information with respect to the experimental hypothesis of interest.

We demonstrate such benefit using the same visual-attention MEG data, to detect the origins of narrow-band oscillations (Fig 2a). The corresponding quality function *f*_*narrow*_ quantifies the ratio of signal power in a frequency range of interest with respect to the total power of the broadband signal. Such a quality function highlights signals with a peaky spectral profile (16), which is of specific interest when studying stimulus-independent ongoing oscillations. We used the data of the ongoing visual stimulus period ([1, 3] s after stimulus onset) to investigate the anatomical origins of three frequency bands of interest: theta (4-8 Hz), alpha (8-13 Hz) and beta (13-30 Hz). The reference broadband signal against which to contrast possible effects in the narrow frequency bands of interest was taken between 2 and 100 Hz.

We compared the *subcorr* statistical maps with the minimum-norm maps of *f*_*narrow*_ (Fig 6). The log-transform of the ratios was not applied because negative effects were of no interest to the question, thus a symmetric measure was not required. A threshold 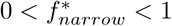 for selecting relevant signal subspace patterns was computed with the bootstrap procedure described in Materials and Methods. In the alpha and beta bands the results were similar between our approach and standard imaging. Commonly observed brain regions as strong sources of these ongoing rhythms were found (see e.g., 17). Alpha activity was prominent in medial occipital-parietal regions; beta activity was stronger over bilateral sensory-motor regions. Alpha band oscillations were also found prominently over the right postcentral region, which parallels the finding of enhanced alpha and beta power during the stimulus period in the same brain area, as shown in the previous section.

**Fig 6.**
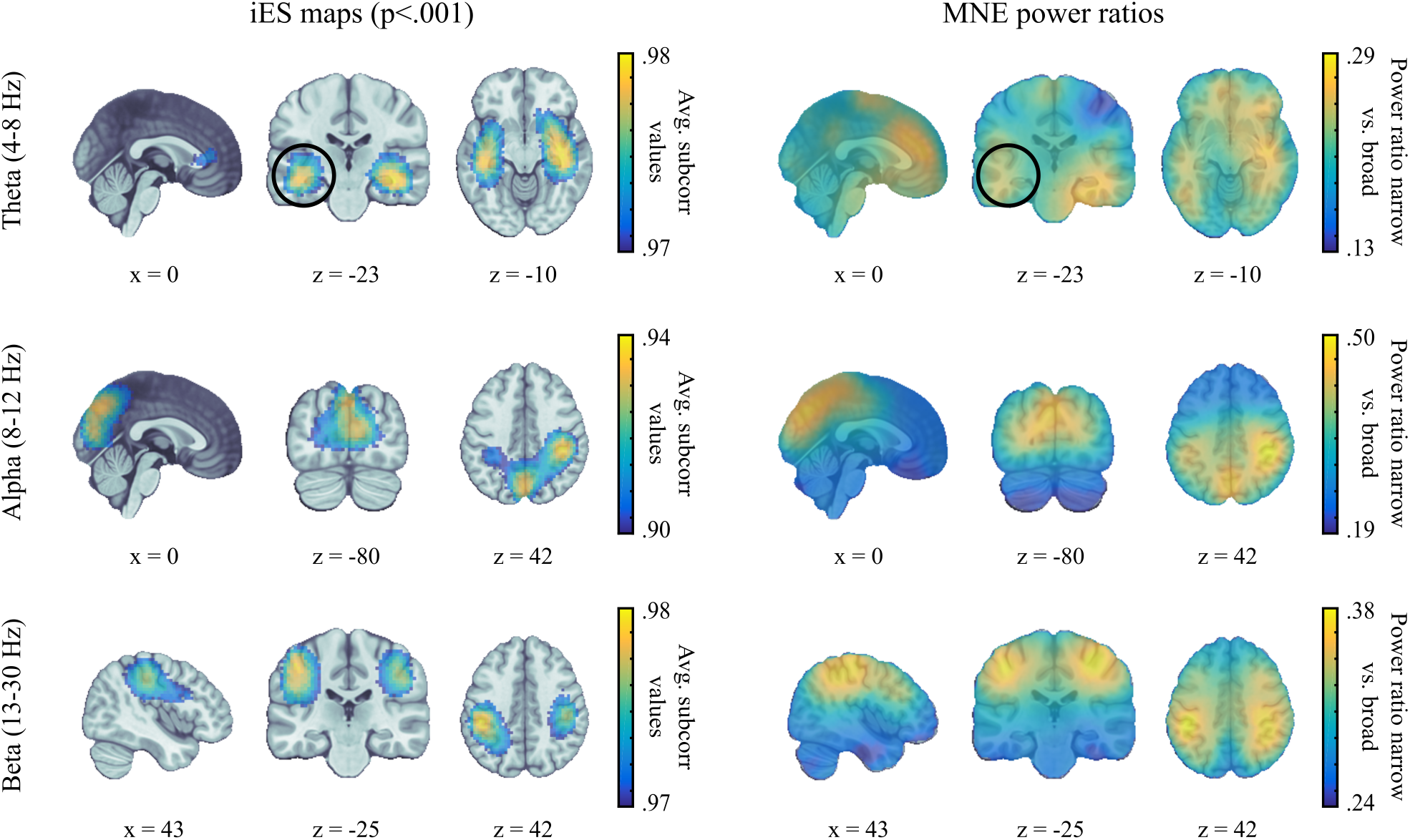
Mapping of narrow-band oscillations. The sources of narrow-band signals were mapped for the theta, alpha and beta frequency bands using *a) iES subspace scanning* and *b) power ratios from minimum norm imaging (narrow-band over broadband 2-100 Hz)*. iES allows for statistical thresholding across the group using permutation procedures that are equivalent for all use cases. The theta band results showed marked differences between the two approaches in deeper, medial temporal regions. iES revealed bilateral sources whereas MNE power ratio maps pointed at predominant source activity in the right hemisphere.

We found differences between iES and minimum-norm maps in the theta band. The *subcorr* statistical map revealed involvement of the medial temporal lobes (MTL) bilaterally, and of medial frontal/anterior cingulate regions. Theta oscillations in MTL, including the hippocampus and parahippocampal regions, have been extensively described (18). Due to their relative depth and therefore lower MEG signal-to-noise ratios (SNR), they have been considered more challenging to detect (19; 20; 21). The MNE power-ratio maps though showed a lateralized distribution of theta activity in the right MTL. We argue that both results are not mutually exclusive: they indicate that both the left and right MTL were consistent sources of theta oscillations in the tested group. However, the effect strength in the right MTL was higher in the average power ratios of theta. Such insight could not be gained with either approach taken separately and required the direct comparison of the iES and MNE statistical maps.

Fig 7 shows simulation results to illustrate and underline further the difference in sensitivity between the iES and standard approach. For each simulation run (300 iterations) we generated five minutes of data. Source time-series with a 1/f spectral profile were generated for 68 source locations distributed evenly across the brain according to the Desikan-Killiany atlas from Freesurfer (22). For two of these locations (precentral left and right), we selectively amplified power in the frequency of interest (8-13 Hz) to obtain a specified ratio *f*_*narrow*_ between narrow-band and broadband power (1-100 Hz). Whereas an *f*_*narrow*_ of 0.6 was targeted for source 1, the targeted *f*_*narrow*_ for source 2 was varied between 0.2 and 0.6. After generating MEG data from this simulation setup, we applied iES (with a *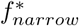*threshold of 0.22) and the standard approach to detect narrow-band oscillations in the frequency band of interest and computed a metric that quantified the probability of detecting both sources of narrow-band oscillations. Fig 7d) shows that the two methods differ systematically: the sensitivity of the standard approach scales with the differences in *f*_*narrow*_ between the two sources, whereas iES’ sensitivity is not influenced by uneven source activity and detects sources above the chosen threshold with a constant probability. This encourages using the different sensitivity profiles of the two methods in conjuncton, to obtain complementary information as shown in the data example above.

**Fig 7.**
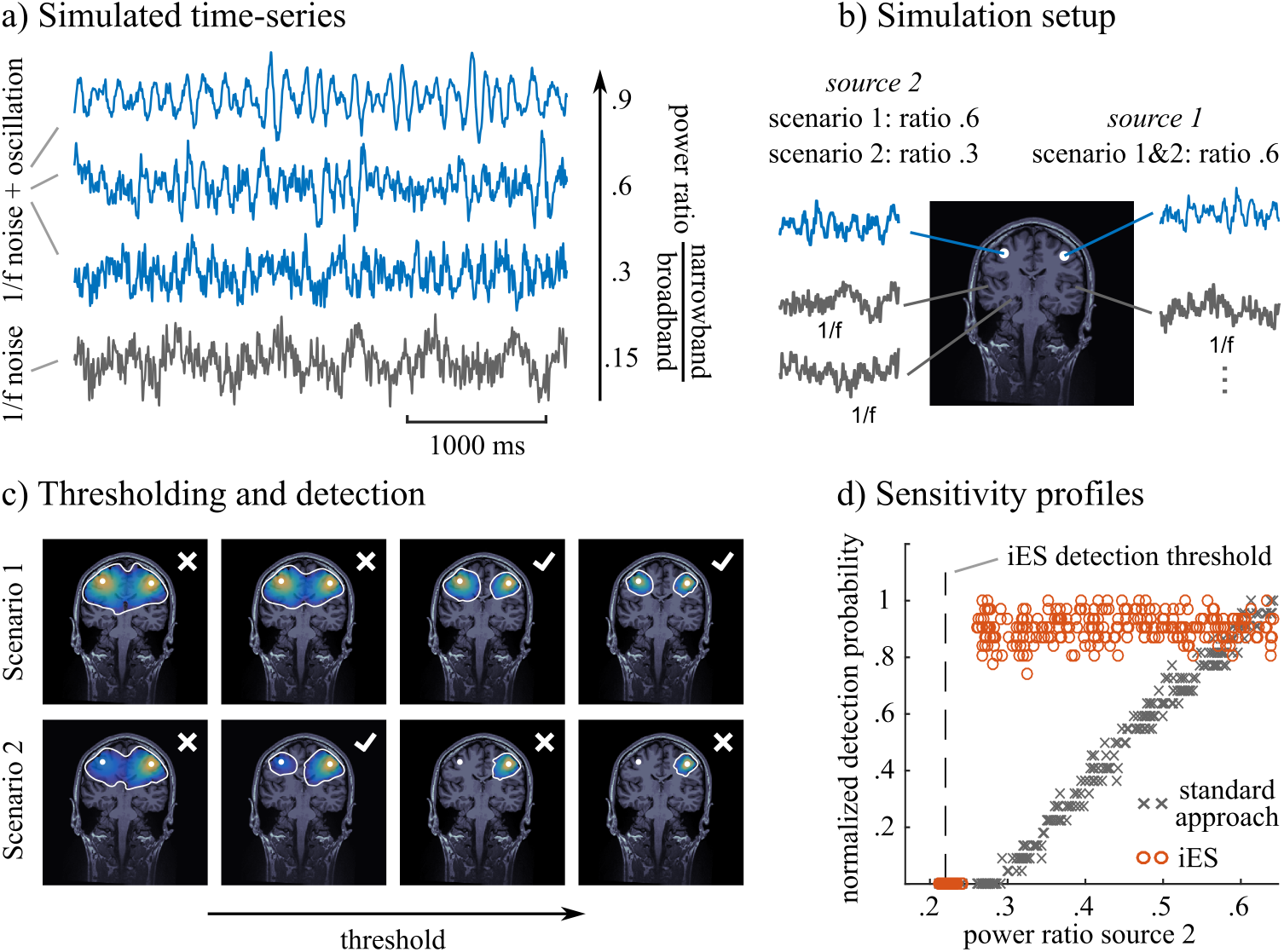
Simulation results comparing sensitivity of iES and standard approach. *a)* Examples of simulated time-series that follow a 1/f spectral distribution (grey trace) or target a pre-specified *f*_*narrow*_, which is the ratio between narrow-band and broadband power (blue traces). *b)* Simulation setup: Two sources of interest in blue targeting pre-specified *f*_*narrow*_ (blue traces) are embedded in background brain noise composed of 1/f signals evenly distributed across 66 locations. *c)* Metric of detection probability: We quantified the probability that the two sources of interest were detected in a source map by using a range of different thresholds: the two sources were detected, if they were contained in two separate clusters after thresholding. Here we show 4 different thresholds in two simulation scenarios using the standard imaging approach. In the first scenario, sources were detected with 2 out of 4 (detection probability: 0.5) threshold values. In the second scenario, sources were detected only with 1 out 4 (detection probability: 0.25) threshold values. This configuration illustrates the issue of concurrent sources with different strengths on the detection of separate clusters of activity. *d)* Comparison of methods: the maps from each simulation run were thresholded using 50 different values to estimate a detection probability as in c). Since the range of data values for both MNE and iES were different, we normalized the detection probability by the maximum value obtained in each method. Thus we did not compare the absolute detection probability between the two methods, but rather how it varied with respect to the difference in *f*_*narrow*_, between the sources of interest.

### Assessment of functional connectivity

Because of spatial smearing, the study of functional connectivity is a challenging problem for MEG and EEG source imaging (see Fig 1). Since the seed region is maximally correlated with itself and neighbouring regions, with correlated time series due to field spread of the MEG/EEG inverse operator, functional connectivity maps tend to be biased towards artificially inflated values of connectivity measures. This issue is discussed in (23) and generally addressed with methods that discard all contributions of zero phase-lag time series, either by orthogonalizing signals (23) or via measures of the imaginary part of coherence (24). However, zero-lag coherence between distant regions is plausible theoretically (25) and was observed physiologically (26). We demonstrate the relevance of iES to address this issue, by studying amplitude correlations in the alpha band (8-13 Hz) with respect to an anatomical seed placed in the sensorimotor cortex. The tested hypothesis was to reveal amplitude correlations with homologous contralateral brain regions (23; 27; 28).

Fig 8 shows results from resting-state data obtained during the same recording session as the visual stimulus experiment. a) shows example time series of co-occurring oscillatory bursts, which form the basis of amplitude correlations between two distant brain regions. The time series were extracted from bilateral central regions. We show occurrences of 270*°/*90*°* phase differences during alpha bursts – note that the phase estimation of MEG source signals has a 180*°* ambiguity due to arbitrary conventions on source direction (29) – and of 180*°/*0*°* phase differences, which would be discarded by other methods (23; 24). We argue that the zero-lag correlations shown here are not spurious, as evidenced by their differences in waveform and amplitude dynamics. This data example provides a proof of principle that studying zero-lag connectivity using MEG is achievable. We next proceeded to map significant inter-regional amplitude correlations in the presence of field spread.

**Fig 8.**
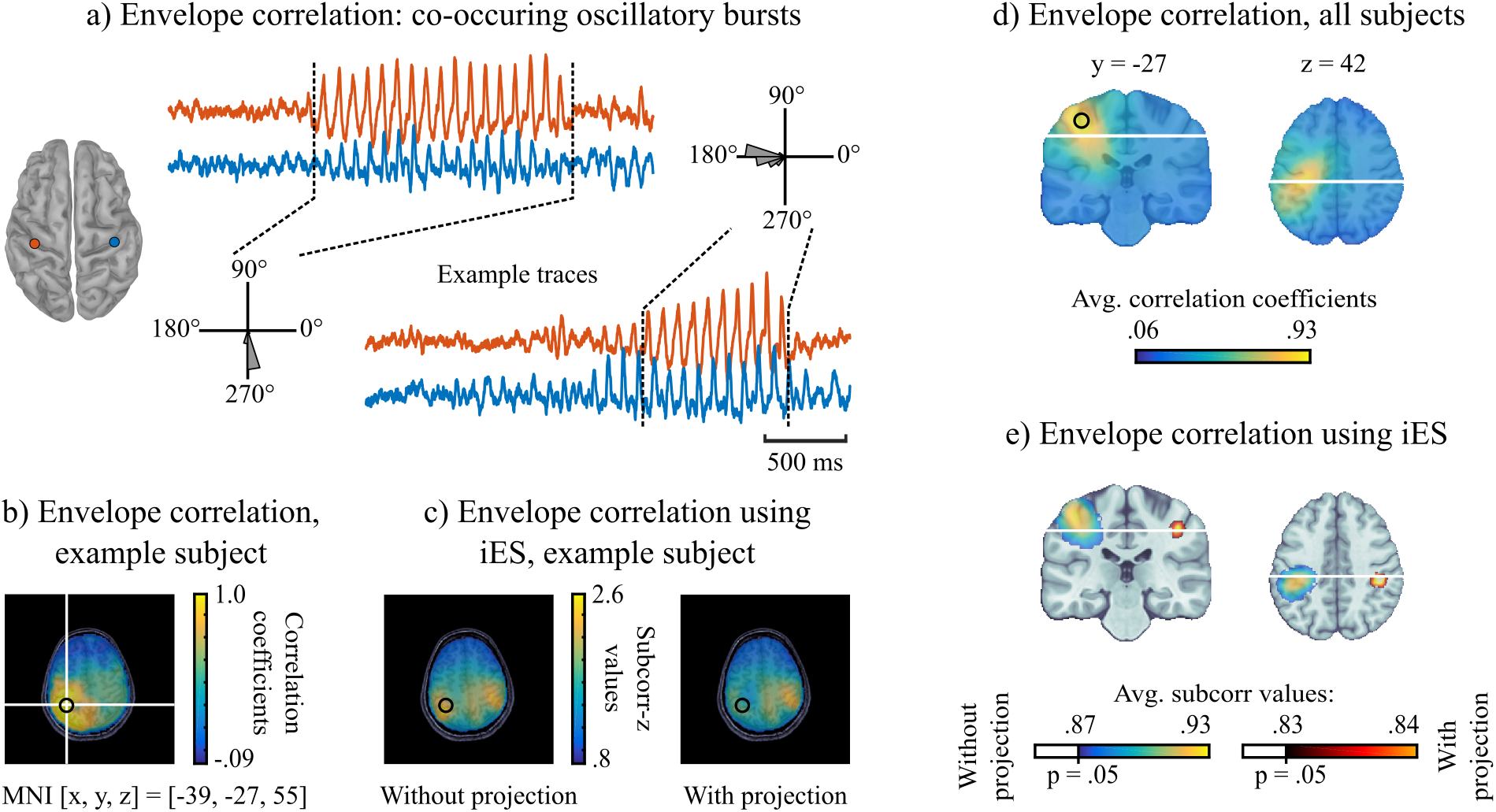
iES mapping amplitude correlations of a seed region with the rest of the brain during rest. *a)* We show example traces of co-occurring oscillatory bursts in the alpha band (8-13 Hz) in the resting-state, from the same MEG sessions as presented in previous sections. The two examples have different phase lags, around 270^*?*^ and 180*°* respectively (which would be discarded in other approaches, see e.g., 23). *b)* Correlation of alpha amplitudes between the seed region (circle) and the rest of the brain, using minimum-norm imaging in an example subject. *c)* (left) iES *subcorr* map showing source locations whose amplitudes correlated with the seed region’s at *r >* .4. The homologous contralateral region is emphasized in this map. (right) the same map with the data projected away from the spatial pattern of the seed region. *d)* Same as b) but averaged over the group *e)* same as c) but averaged over the group and statistically thresholded using the permutation approach explained above.

We extracted the source time series **y**_*ref*_ from the left central sulcus location that was the closest to the activation peak (MNI coordinates [-39, -27, 55] mm) corresponding to the search term ‘finger’ in the Neurosynth meta-analysis tool (30). We defined the iES quality function *f*_*ampcorr*_(**s**, **y**_*ref*_) as the correlation between the amplitude of other source time series **s** and **y**_*ref*_. The outcome to this optimization process is the third use case of iES and is illustrated in Fig 2a. The optimization is done using a solution described in (31). We set a correlation threshold of *r >* 0.4 for spatial components to be included in the signal subspace for the *subcorr* analysis.

Fig 2b-c shows – in a single subject example – that iES was able to reveal the contralateral anatomically-homologous region as the primary distant connected region with the reference brain location. The conventional minimum-norm based map of correlation values was dominated by spurious crosstalk correlation surrounding the seed region. The performance of iES is explained by the equalized contribution of spatial components that are consistent with the embedded hypothesis (*r >* 0.4). To further limit the contribution of the seed region to the data, it is possible to project the signal subspace and forward fields away from the spatial forward field of the seed region, as illustrated in Fig 2c. The group analysis further reveals that connectivity maps were dominated by crosstalk effects from the seed reference signal, both in the minimum-norm based maps and in the raw *subcorr* map (Fig 2d-e). Projecting away the seed’s contribution before computing every subject’s maps was necessary to confirm the hypothesized contralateral coupling. Note that with iES and in contrast with other approaches (23), the temporal dynamics of the seed region are not projected away from the data; only the spatial topography of the seed region is subtracted from the sensor data. Thus iES does not exclude the detection of physiological zero-lag coupling.

### Applicability to a wide range of experimental questions

As summarized in Fig 2a, iES can be used for a greater variety of experimental designs: whenever a reference signal **y**_*ref*_ defined 1) on a trial-by-trial basis or 2) as a continuous signal is considered, *f*_*ampcorr*_ is used to obtain *subcorr* maps of sources, whose source dynamics correlate with **y**_*ref*_. We illustrate such case in Fig 9, using simultaneous MEG and pupil diameter recordings.

**Fig 9.**
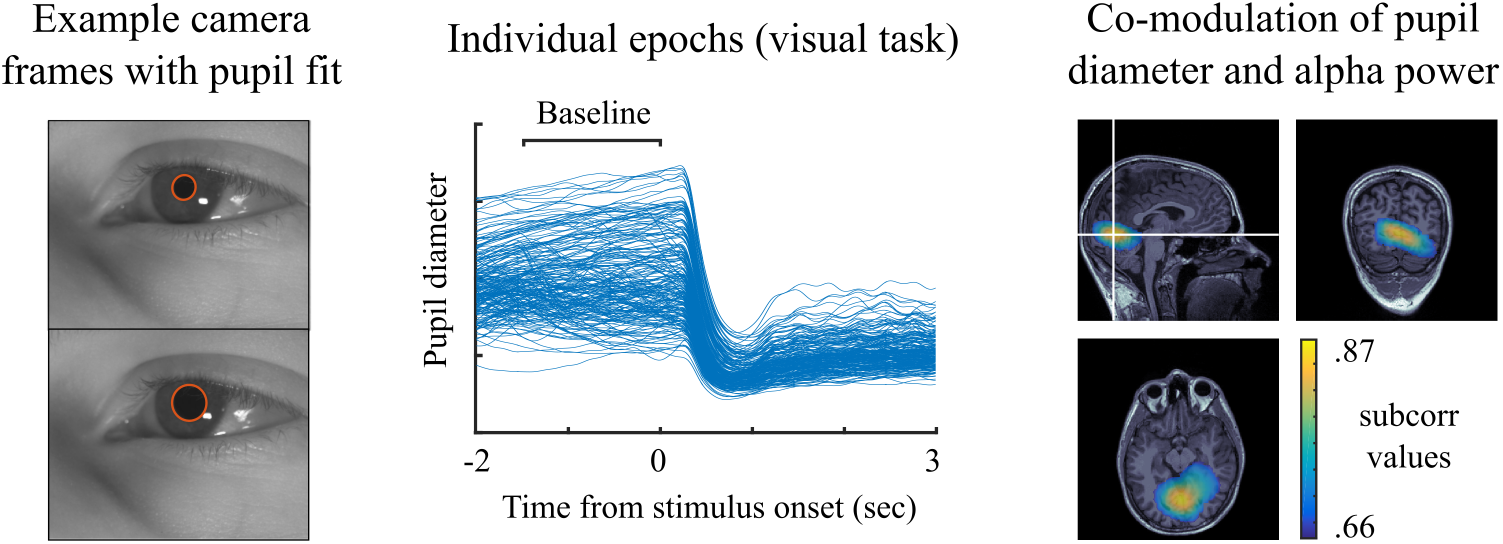
Mapping correlation of a peripheral signal with neural oscillation amplitudes. *a)* video frames from an eye tracker camera during a MEG recording at rest, pupil diameter was extracted using a fitted ellipse. *b)* pupil diameter time-series time-locked to a visual stimulus onset (overlaid trials). The gray bar indicates the baseline time period for analysis. *c) subcorr* map showing sources whose amplitudes correlate with pupil diameter across trials during the visual task. The signal subspace for this analysis contained one significant component which signal correlated at *r* =0.44 with changes in pupil diameter. The maps were threshold at 75% of the maximum value.)

We first formed the iES hypothesis based on recent demonstrations in mice (32) that continuous pupil diameter fluctuations correlated with alpha power at rest. Measures of pupil diameter were extracted from continuous video eye-tracking recordings by fitting an ellipse to the pupil on a frame-by-frame basis. We demonstrate the case of trial-by-trial correlations by analyzing pupil diameter changes prior to visual stimulus presentation from data presented in Fig 3. The signal subspace was defined with spatial components who were deemed significant below a p-value of 0.05 computed by a shuffling procedure across trials. The iES maps indicated brain regions in the occipital cortex, consistent with the upcoming onset of a visual stimulus.

We emphasize that a specific strength of the iES approach is its versatility: it can be extended to a great variety of experimental designs and research hypotheses, since the experimental question is formulated as an optimization problem. We derive in Methods the mathematical formulation for iES coherence with a reference signal, as an additional experimental use case. The experimental hypotheses discussed here all have corresponding quality functions that can be solved analytically. An identical framework can be used for hypotheses that require numerical optimization of the corresponding spatial filters. We foresee that the introduction of the iES approach will establish a generic framework for an increasing number of experimental contexts related to a growing diversity of research questions.

## Materials and Methods

### MEG data

#### Participants

17 healthy participants were recruited (21-45 years; 5 female). The study was approved by the Montreal Neurological Institute’s ethics committee, in accordance with the Declaration of Helsinki. All participants gave written informed consent and were compensated for their participation.

#### Stimuli

Subjects were presented with a variation of the visual stimulation paradigm in (9): A circular sine wave grating (diameter of 5°with 100% contrast) contracts towards the fixation point (velocity: 1.6 deg/s). The contraction accelerated (velocity step to 2.2 deg/s) at an unpredictable moment between 3-5 seconds after stimulus onset. Subjects had to indicate with an index-finger button press that they detected the velocity change. The button press ended one trial and the stimulus was turned off. Inter-trial intervals were 5 seconds long with a jitter. During the inter-trial interval subjects were presented with a central fixation cross. Stimuli were generated using the Psychophysics Toolbox (33).

#### Experimental Procedure

Participants received both oral and written instructions on the experimental procedure and the task. The recording session started with a 5-minute resting state run with eyes open. The participants were presented with 10 test trials, to become familiar with the task. They were then presented with a total of 240 stimulus sequences (trials). Participants performed 60 trials per acquisition block. After each block, they received a feedback on the accuracy of their responses. A trial was considered correct if subjects responded within 500 ms after the actual velocity change occurred. Between each block, the participants were given a break of self-determined length. After completion of the 240 trials, subjects were given a 15-minute break. A further 5-minute resting-state recording concluded the session.

Fig 3, 4, 5 and 6 are based the visual experiment phase of the recording sessions, whereas figure 8 is based on the two resting-state sessions. Fig 9 is based on the visual experiment phase of one subject, during which eye videos were recorded concurrently with MEG. For this we used a Point Grey Flea3 camera, capturing at 15 frames per second, with an infrared filter and infrared illumination of the subject’s face to improve the contrast for pupil segmentation.

#### Data acquisition

The participants were measured in a seated position using a 275-channel VSM/CTF MEG system with a sampling rate of 2400 Hz (no high-pass filter, 660 Hz anti-aliasing online low-pass filter). Three head positioning coils were attached to fiducial anatomical locations (nasion, left/right pre-auricular points) to track head movements during recordings. Head shape and the locations of head position coils were digitized (Polhemus Isotrak, Polhemus Inc., VT, USA) prior to MEG data collection, for co-registration of MEG channel locations with anatomical T1-weighted MRI. Eye movements and blinks were recorded using 2 bipolar electro-oculographic (EOG) channels. EOG leads were placed above and below one eye (vertical channel); the second channel was placed laterally to the two eyes (horizontal channel). Heart activity was recorded with one channel (ECG), with electrical reference at the opposite clavicle.

A T1-weighted MRI of the brain (1.5 T, 240 x 240 mm field of view, 1 mm isotropic, sagittal orientation) was obtained from each participant, either at least one month before the MEG session or after the session. For subsequent source analyses, the nasion and the left and right pre-auricular points were first marked manually in each participant’s MRI volume. These were used as an initial starting point for registration of the MEG activity to the structural T1 image. An iterative closest point rigid-body registration method implemented in Brainstorm (15) improved the anatomical alignment using the additional scalp points. The registration was visually verified and adjusted manually, if necessary.

### MEG data preprocessing

All MEG data analysis steps were performed with Brainstorm (15), with the novel approaches described in this paper implemented as a Brainstorm plug-in written in MATLAB (available through: https://github.com/pwdonh/ies_toolbox).

#### Artifact removal and rejection

Eye-blink and heart-beat artifacts were removed from MEG data using a PCA-based signal source projection (SSP) method, using recommended procedures (34). The ECG and EOG channels were used to automatically detect artifact events. Noisy MEG channels were identified by visually inspecting their power spectrum and removing those who showed excessive power across a broad band of frequencies. The raw data were further visually inspected to detect time segments with excessive noise e.g., from jaw clenching or eye saccades. Sinusoid removal at the power line frequency and harmonics (60, 120, 180 Hz) was applied to the continuous data. A high-pass filter above 1 Hz was also applied to reduce slow sensor drifts. The MEG data were centered around the baseline mean after epoching. All the filters used in the current study are zero phase shift non-causal finite impulse filters coded and documented in Brainstorm.

MEG data were epoched to the interval [-2, 3] seconds around the visual stimulus onset. We refer here to the stimulus period as the interval [1, 3] seconds, and the baseline period as [-2, 0] seconds with respect to visual stimulus onset.

#### Intra-subject coregistration

Prior to the computation of signal subspaces, we performed a between-run coregistration of the MEG data based on recorded head positions, using the movement correction method similar to (35) available in Brainstorm. Briefly, we computed forward models **G**_*k*_ based on the head positions of different runs *k* and **G**_*avg*_ using the average head position. Then we computed coregistration operators **Ω**_*k*_ to project the MEG data from different runs into the same space as **Ω_k_X_k_**. **Ω**_*k*_ was computed as **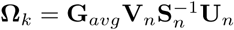** using the singular value decomposition (SVD) **G**_*k*_ = **USV** truncated corresponding to the largest *n* singular values. The index *n* is set so as to preserve 99.99% of the squared singular value spectrum. We additionally took into account that different runs had slightly different SSP projectors applied (see above). We thus apply these projectors to the forward fields of individual runs before computing the coregistration operators.

#### Source models

We defined a volumetric source grid on the MNI152 2009c nonlinear anatomical template (36), using an adaptive procedure: an outer layer of 4000 grid points was produced based on a brain envelope covering cortical and subcortical structures. This outer layer was then shrunk and downsampled by a factor of 2.2. This procedure was repeated to result in a total number of 20 layers containing 25740 grid points. For each subject, we computed a linear transform of individual anatomy to MNI coordinates using affine coregistration as implemented in SPM12 (function spm maff, 37). We applied the inverse linear transform to project the default source grid onto each subjects’ individual anatomy.

Forward modeling of neural magnetic fields was performed using the overlapping-sphere model (38). Conventional MEG source imaging was obtained by linearly applying the weighted-minimum norm operator, with default settings in Brainstorm (2). The weighted-minimum norm operator included an empirical estimate of the noise covariance at the MEG sensor array, obtained from empty-room recordings (34).

For the extraction of a seed time-series for functional connectivity analysis in Fig 8 we found the optimal dipole direction at the seed location using SVD of the filtered 3 component time-series extracted with the MNE kernel *K*(*ρ*).

### iES formulation

The iES method described in this paper is based on subspace scanning, which processes the entire spatio-temporal MEG data matrix **X**, instead of reconstructing neural activity independently at each time point (see e.g. MUSIC, 6). The method features two steps, as shown in Fig 2: 1) extraction of the relevant spatial patterns from the data (signal subspace identification), and 2) scanning of the source space for contributions that explain the identified spatial patterns (subspace scanning step per se).

#### Subspace scanning

First a set of MEG topographies is identified that captures the signal components of the MEG data matrix. We define the notion of *signal* in the following section. In (6), this first step was equivalent to performing a PCA of the event-related average MEG signals. The *D* components corresponding to the largest PCA eigenvalues spanned the signal subspace: span(**P**^*s*^). A particular MEG topography **v** lies within the signal subspace, if there exists a linear combination **t** of the columns of **P**^*s*^ such that **v** = **t**^*T*^ **P**^*s*^. A geometric measure quantifies how close a particular MEG topography lies to the signal subspace. For instance, the cosine of the angle between **v** and the projection of **v** onto **P**^*s*^ is a suitable measure (6).

A dipolar source at location *ρ* is described by an orientation (*θ*) and an amplitude (*a*) parameter. All the MEG topographies that can be generated by this source are described by a linear combination of the forward fields of dipoles along the three orthogonal spatial directions (2). The resulting 3-column forward field matrix **G**(*ρ*) thus also spans a subspace. A MEG topography produced by this dipole lies within the signal subspace if there exist linear combinations **u** and **t** such that **u**^*T*^ **G**(_*ρ*_) = **t**^*T*^ **P**^*s*^. Due to noise in measurements and inevitable approximations in the forward model metric, a perfect match cannot be expected. Thus we use the subspace correlation metric as the cosine of the smallest principal angle between subspaces

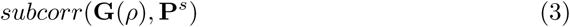

as defined in (6). This metric quantifies how close the two subspaces lie to each other, and thus how well a dipole source at the scanned location fits the signal subspace. This metric is applied at each possible location across the anatomical volume.

#### Computing the signal subspace

The signal subspace is defined by the span of the full column-rank *M×D* matrix **P**^*s*^ where *M* is the number of sensors and *D* is the dimensionality of the subspace. Thus **P**^*s*^ is a collection of *D* non-collinear sensor topographies. In the standard MUSIC case, the first column of the signal subspace **p**_1_ is a vector/topography that, when applied to the averaged MEG data **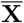**, results in a signal that has maximum variance (broadband power): it is a solution to the optimization problem

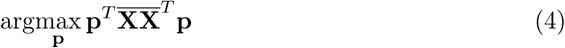

subject to a norm constraint on **p**. The next subspace column **p**_2_ is the solution to the optimization problem

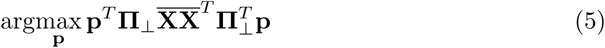

where **π**_⊥_ is the orthogonal projector away from the first subspace column **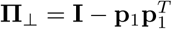**. This corresponds essentially to a PCA of the event-related average **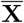**. More generally, the signal subspace could be constructed by the solution of an optimization problem

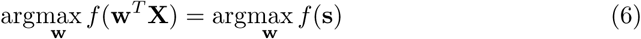

where the function *f* is chosen according to the experimental question of interest. The time-series **s** is the signal obtained by applying the spatial filter to the data as **s** = **w**^*T*^ **X**. In standard MUSIC, the experimental question of interest is to find the sources that have the strongest contribution to the event-related responses, thus the quality function is

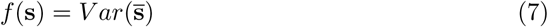

where **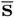** is the event-related trial average of s. However, many other experimental questions can be expressed as an optimization problem. For example we might be interested in finding sources whose power is correlated with a reference signal **y** (such as an EMG recording or an audio stimulus envelope). In that case we would set the quality function as

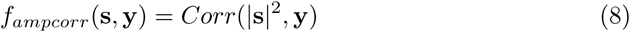

where *Corr*(**a**, **b**) is the correlation of signals **a** and **b**. While, in principle, it is possible to use any quality function and proceed with numerical optimization, the subspace method is specifically attractive for quality functions that can be solved analytically for computationally efficient implementations. Here we focus on a set of quality functions that can be solved using the generalized eigenvalue problem (GEP). We show solutions for four different experimental use cases, three of which are illustrated in Fig 2a).

##### Subspace computation using the GEP

The generalized eigenvalue problem (GEP, see e.g. 39) **Aw** = *λ***Bw**, for symmetric matrices **A** and **B**, arises in optimization situations like

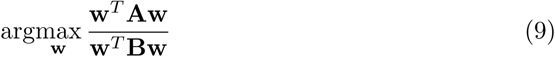

or equivalently

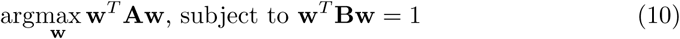

To show how the GEP can be used to define a subspace, we focus on **a) induced responses** as a first use case (see Fig 2a). Here we are interested in finding sources whose power in a frequency band of interest [*f*_1_, *f*_2_] differs between two conditions or time periods, e.g. stimulus and baseline periods. Thus the quality function becomes

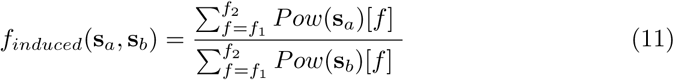

where *Pow*(**s**_**a, b**_)[*f*] is the power of a signal s at frequency *f* in time periods *a* and *b*. The power of a signal in a given frequency band [*f*_1_, *f*_2_] can be approximated by the variance of the signal filtered in that frequency band. The quality function can thus also be written as

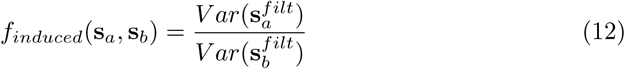

where superscript *filt* indicates that the signal was filtered in the frequency band of interest. For readability, we will drop this superscript in the following. Since the bandpass-filtered signal is zero mean, we can compute the variance using the dot product 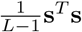 where L is the number of time samples. The quality function thus becomes

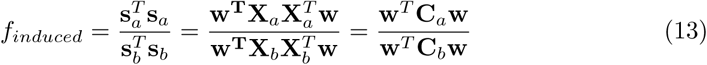

where **C** is the covariance matrix of the filtered MEG signals **X** and **w** is a spatial filter topography. This is now in the form of the GEP shown above and has been used in the field of brain-computer interfaces as Common Spatial Patterns (CSP, e.g. 3; 40). Alternatively, one can define the quality function directly in the frequency domain, and compute **C**_(*a,b*)_ as the average of the real part of the cross-spectral density matrices in the frequency band of interest

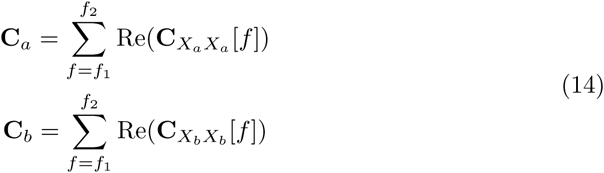

where **C**_*XX*_ [*f*] is the *M × M* MEG cross-spectral density matrix at frequency *f*.

The GEP can now be solved by defining a whitening projector

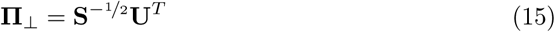

from the SVD: **USV**^*T*^ = **C**_*b*_, which equalizes the variance along the principal axes of **C_b_**, as required in the constraint of Eq 10. We then solve the ordinary eigenvalue problem

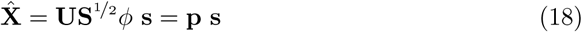

where the eigenvector *ϕ* is now a spatial filter in the whitened data space. The eigenvalue *λ* provides the ratio of power in the two conditions, thus is equal to *f*_*induced*_. This means that the signals of interest, which maximizes the quality function in Eq 12, can be estimated from the MEG data as

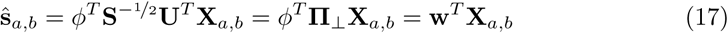

where **w** combines the two steps of whitening (**Π**_**⊥**_) and filtering in whitened space (*ϕ*^*T*^) to obtain a spatial filter in the data space as in Eq 13. The data generated by a specific source signal can in turn be estimated by

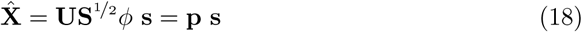

where **p** is the spatial pattern vector, or forward field, of the source signal in sensor space, since an inverse whitening step (**US**1*/*2) is applied to the forward pattern in whitened space (*ϕ*). (see 41, for further discussion on the distinction between spatial patterns and filters).

Solving the GEP this way, one obtains *M* spatial patterns **p**_*j*_ that can be ordered according to their corresponding quality function scores *f*_*induced*_ = *λ*. The columns of the signal subspace matrix **P**^*s*^ are then defined by the spatial patterns that exceed a threshold *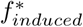*, yielding an *M* by *D* subspace matrix, where *D* is the number of spatial patterns exceeding the threshold. We discuss the estimation of this threshold below. As described above, the anatomical source space can then be scanned by computing subspace correlations with the forward fields at each source location, using

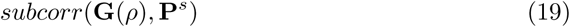

##### Additional use cases based on the GEP

We describe three other use cases that can be solved using an appropriate quality function in combination with the GEP (see Fig 2a). The solutions are convenient in that they only require to change the definition of **C**_*a*_ and **C**_*b*_ in Eq 13.

We define a signal showing **b) narrowband oscillations** as a signal that has increased relative power in a frequency band of interest with respect to broadband power. We thus define a signal frequency band of interest 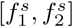 and a broad noise frequency band 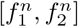. The quality function then becomes

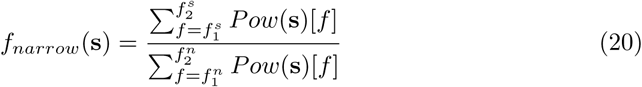

Analogously to Eq 13, this quality function can be expressed in the form of the GEP as

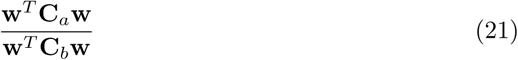

where

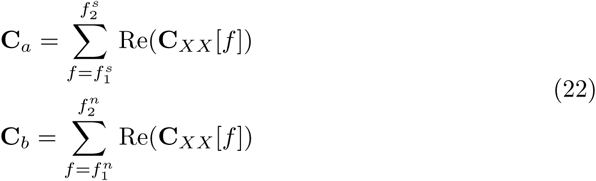

Solving the GEP, we obtain spatially filtered signals that are ordered according to their ratios of power *f*_*narrow*_ in the signal and noise frequency bands. This approach is similar to what has been described in (16) as spatio-spectral decomposition.

As next use case, we consider the case of **c) amplitude modulation** using a solution described in (31). Here we wish to find sources whose amplitude fluctuations in a frequency band of interest covary with the value of a reference variable. This might be a slow time-varying signal **y**_*ref*_, or a variable that is defined on a trial-by-trial basis such as reaction time or task difficulty. Here we describe the former case, but the latter follows easily (31).

The data are split into epochs denoted by the index *e*. Epoch length needs to be short enough to allow capturing fluctuations in the reference signal **y**_*ref*_, and long enough to estimate the power of data signals **X** filtered in the frequency band of interest. The quality function can then be expressed as

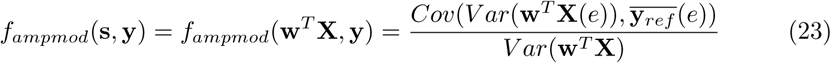

where **X**(*e*) denotes the data matrix in epoch *e*. We thus maximize the covariance between the power of **s** = **w**^*T*^ **X**(*e*) and the value of **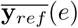** normalized by the power of **s** = **w**^*T*^ **X** = 1. Assuming that **y**_*ref*_ is a zero-mean and unit-variance signal, this can be solved in the GEP framework by setting

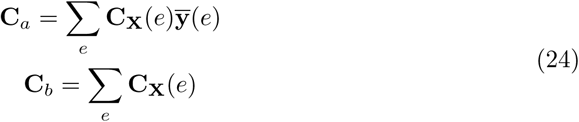

where **C_X_**(*e*) is the covariance matrix of the filtered MEG signals in epoch *e*. **C**_*a*_ thus represents a weighted (by 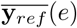 (*e*)) average, and **C**_*b*_ the unweighted average of the single epoch covariance matrices. Please refer to (31) for a derivation of these results. We can obtain the correlation values from the above as

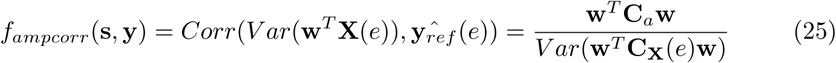

In the analysis examples we used *f*_*ampmod*_ to compute the spatial filter basis using the GEP in a computationally efficient manner. Ordering and selecting the components to be included in the signal subspace was then based on *f*_*ampcorr*_.

When the research hypothesis requires testing for source dynamics that are **d) coherent with a reference signal y**_*ref*_ at a specific frequency, the quality function becomes

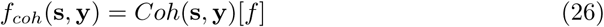

The reference signal **y**_*ref*_ can be an external stimulus such as the envelope of an audio signal, a simultaneously measured peripheral signal such as EMG, or a neural time-series extracted using source imaging.

Magnitude squared coherence is computed as the ratio of cross-spectral to auto-spectral densities as

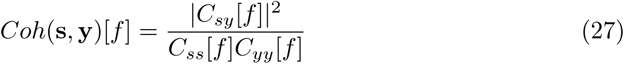

where *C*_*sy*_[*f*] is the cross-spectral density between signals **s** and **y** at frequency *f*, and *C*_*yy*_[*f*] is the auto-spectral density of signal **y**. Since *C*_*yy*_[*f*] is constant, we can leave it out of the quality function and remain with

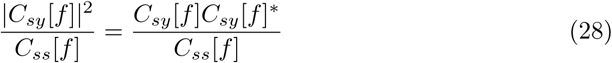

Now setting **s** = **w**^*T*^ **X** we get

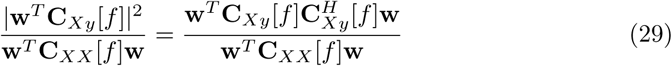

where **C**_*Xy*_[*f*] is the column vector containing the cross-spectral densities between the reference signal and the MEG signals, and the *H* superscript stands for conjugate transpose. The optimization problem can be solved by invoking the GEP as in Eq 21 and setting

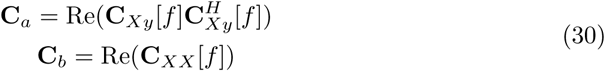

Note that, as the matrix **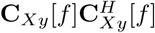**is Hermitian, **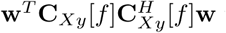** will be a real number and thus be equal to 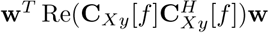. Hence we only need to keep the real part for **C**_*a*_.

##### Covariance regularization

As described above, at the core of our approach lies the computation of a subspace using the GEP. This requires the inversion of matrix **C**_*b*_, which can be numerically unstable if some of its singular values are small (see Eq 15 for its influence on the whitening projector). We thus regularize matrices **C**_*a*_ and **C**_*b*_ by adding values to the diagonal as

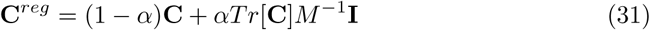

where *α* is a regularization parameter. We refer to this technique as *diagonal loading*. Fig 10 shows the effect of regularization on the power of individual components, illustrated using the data from Fig 3 (induced responses, stimulus vs. baseline). Regularization affects mostly the components with smallest power values, which would have detrimental effects during inversion. Their power ratios *f*_*j*_ after regularization will be very close to

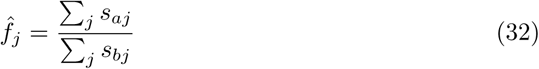

**Fig 10.**
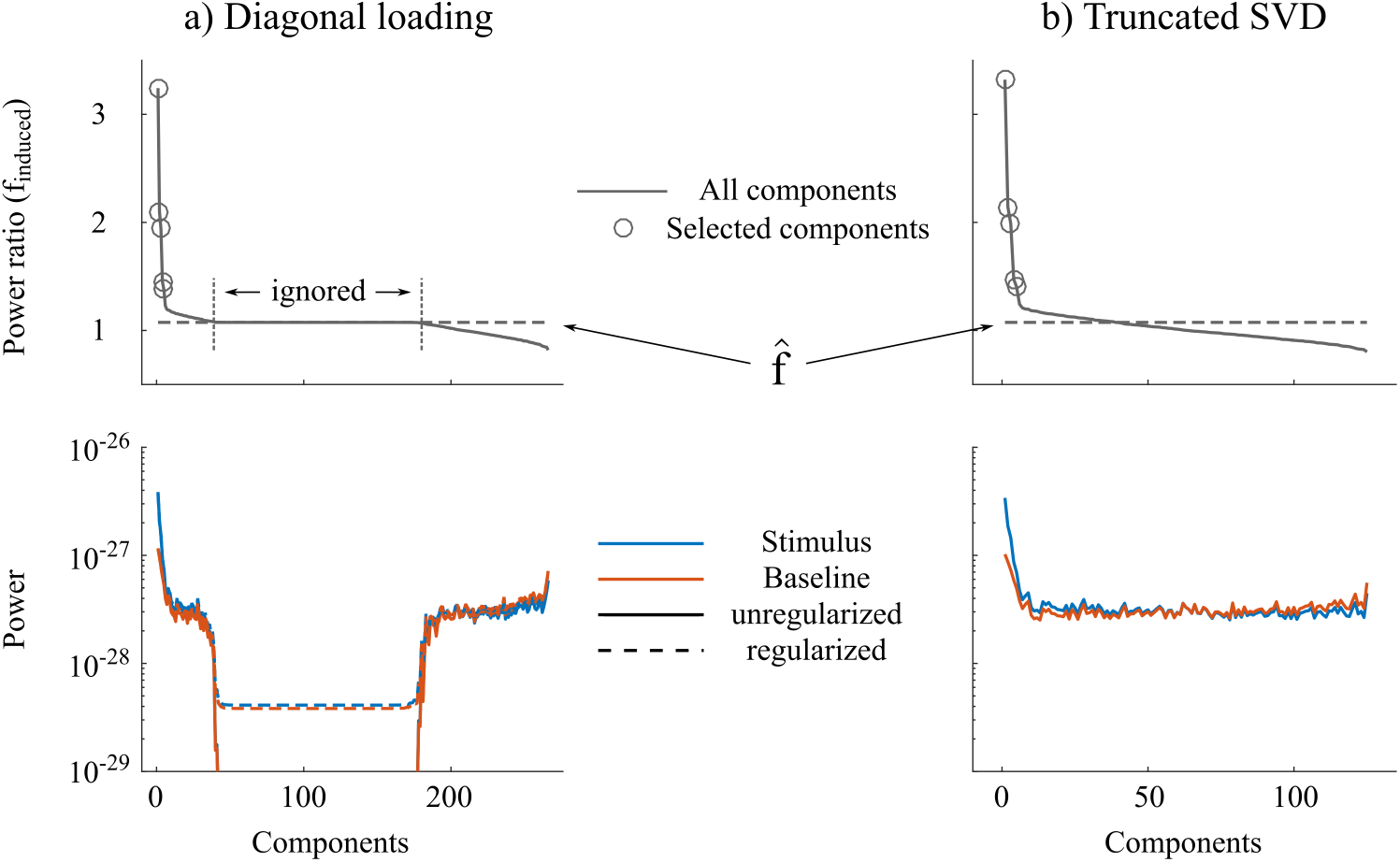
Regularization methods for covariance estimates. For illustration we show GEP results from the single-subject data of Fig 3a comparing gamma power between stimulus and baseline periods. The top panels display the power ratio *f*_*induced*_, the bottom panels display the power associated with each component, computed as **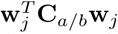** in the baseline and stimulus period. In *a)* component-wise power computed from regularized and unregularized covariance matrices are compared using the *diagonal loading* method. In *b)*, regularized results using the *truncated SVD* method, which results in a smaller number of components.

Where *s*_(*a,b*)*j*_ are the singular values of the narrow-band covariance matrices **C**_(*a,b*)_ computed from stimulus (a) and baseline (b) periods, respectively. This can be understood as the *expected f* based on the overall power across sensors in both time periods. Because these components do not carry physiological information, we make sure to never include them in the signal subspace. They were detected using a simple bootstrap procedure: a confidence interval on the mean power of each component (*s*_*aj*_ and *s*_*bj*_) is estimated from the unregularized data by sampling with replacement from epochs *e* = 1, *…, E*. We then verify whether *s*_*aj*_ and *s*_*bj*_ computed from the regularized data lie within the 99.9% confidence interval, and discard the components where this is not the case. We used this approach in the analysis examples on induced responses and narrow-band oscillations (Fig 3, 4, 5, 6 and 7) and set *α* = 0.05.

We also describe an alternative regularization approach, referred to as *truncated SVD*. This entails removing the columns associated with the smallest singular values from **U** during the computation of the whitening projector in Eq 15. We define the regularization parameter *E* and keep the singular values making up 100(1*-∈*)% of the cumulated singular value spectrum. We see in Fig 10b that this results in a smaller number of components extracted from the GEP, however yielding similar *f* values at both ends of the spectrum. We used this approach in the analysis examples on amplitude modulation (Fig 8 and 9) and set *∈* = 0.001.

##### Estimating the dimensions of the signal subspace

The estimation of the dimension of the signal subspace can be obtained by setting a threshold on the quality function score *f* in a hypothesis-driven way, as a measure of effect size. These scores are readily interpretable as e.g., the power ratio between conditions (induced responses) or the correlation between a reference signal and neural amplitude time-series (amplitude modulation). In the following we show how we can set a threshold based on approaches from non-parametric statistical testing.

For **a) induced responses** we compare oscillatory power between two conditions using permutation testing. We describe the case of stimulus-baseline contrast where, for each epoch *e*, a data matrix for baseline and stimulus periods is available to perform a paired test. Other cases can be derived easily using standard approaches in non-parametric statistics (see e.g. 4; 5). Under the null hypothesis of no difference, the condition labels are exchangeable with respect to the statistic of interest *f* (here *f*_*induced*_), which is the ratio of power between the two conditions

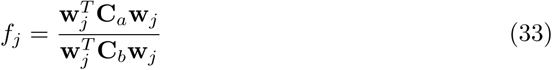

which is defined for each of the potential columns *j* of the signal subspace matrix **P**^*s*^.

The data are divided in *e* = 1, *…, E* epochs, from which we compute empirical covariances **C**_*a*_(*e*) and **C**_*b*_(*e*). We run *O* permutations, where at each iteration, a binary permutation vector *ω* of length *E* is drawn. At each permutation we solve the GEP based on the permuted condition labels and compute a maximum statistic as

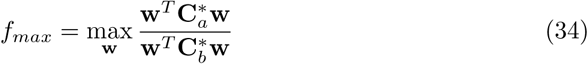

Where

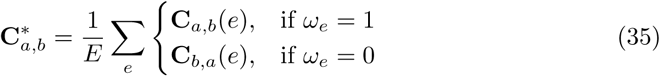

and the *f*_*max*_ values are logged at each iteration. We then obtain a null distribution of *f*_*max*_ (assuming exchangeability of the condition labels) against which to test the observed *f*_*j*_’s to obtain a permutation p-value. In this paper we use *O* = 600 permutations.

In the second use case **b) narrowband oscillations** the power ratios *f*_*j*_ (*f*_*narrowband*_) will differ depending on the frequency band of interest. Due to 1/f in electrophysiology power spectra, low-frequency bands have higher *f*_*j*_’s than high-frequency bands. To find spatial patterns **p**_*j*_ whose relative power stands out from the rest of the activity, we define an expected *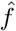* as the ratio of overall power in the narrow- and broad frequency bands

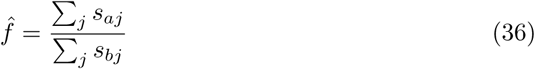

where *s*_(*a,b*)*j*_ are the singular values of the narrow- and broadband cross-spectral densities **C**_(*a,b*)_ as defined in Eq 22. We use a bootstrap procedure to find the dimensions *j* that reliably lie above this expected power ratio. A large number of bootstrap samples can be obtained by sampling with replacement from the epoched data, and logging the mean values over the *f*_*j*_(*e*) of each selected epoch. A confidence interval based on the obtained bootstrap distribution is obtained and a dimensionality *D* up to which the confidence interval does not contain the expected power ratio is therefore defined. In Fig 6, we used a confidence interval of 99.9% to define the signal subspace.

Use cases **c) amplitude correlation** and **d) coherence** compute a measure of temporal association between time-series. The temporal ordering between the reference signal **y** and the data **X** can be scrambled under the null hypothesis of no association. Thus we compute a null distribution of the respective *f* values by randomly assigning epoched data of **y**(*e*) and **X**(*e*). At each iteration the GEP is solved resulting in a single null distribution of *f* ^*\**^ against which to test all the observed *f*_*j*_.

#### Projection of the seed topography for functional connectivity analyses

We can project out the topographic contribution of the seed location *ρ*_*s*_ in a functional connectivity analysis as shown in Fig 8. Using subspace correlation we define the topography **g** at location *ρ*_*s*_ with orientation *θ* that maximizes the fit with the signal subspace as

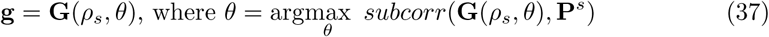

Then we find the orthogonal projector

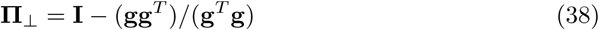

to be applied to both the signal subspace and the leadfield matrices, so that we can scan the source space as

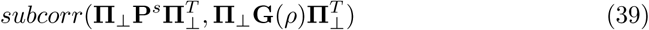

#### Group analysis

##### Testing the effect prevalence

We have derived the dimensionality of each subject’s subspace using tests described in the previous section. The subspace matrix **P**_*i*_ of subjects *i* = 1, *…, N* has an estimated dimensionality of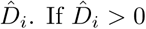, one can claim that subject *i* shows the effect of interest, i.e. there is a spatial dimension in which the null hypothesis can be rejected. For example, the effect of interest could be that the power of gamma oscillations in one spatial dimension of the subject’s sensor data is stronger during presentation of a stimulus than during rest. As a first step for group-level analyzes, we test if the mere presence of the effect is generalizable to the population. If the effect is deemed generalizable, we run a procedure to test if there exist consistent source spatial locations across the group, where the effect originates from (see next section).

The first step requires formulating a prevalence hypothesis (see e.g. 10; 42; 43). In this framework, a true effect is assumed to be present in a proportion *γ* of the population. Hence if a subject *i* is randomly selected from the population

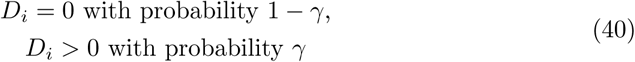

We then specify a prevalence null hypothesis that *γ* is smaller than or equal to a certain proportion *γ*_0_. In order to claim that the effect is generalizable to the population, an intuitive value for *γ*_0_ is 0.5, i.e. the effect would be present in the majority of the population. If we observed that *K* out of *N* subjects showed an effect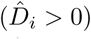, we can define a *p*-value for the likelihood of *K* or more out of *N* subjects showing an effect, if the prevalence across the population is smaller than or equal to *γ*_0_:

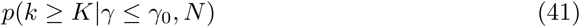

If this p-value is below a specified significance level, the effect is deemed generalizable to the population.

A certain subject *i* can show an effect both if the effect is actually present (with sensitivity *β*), or because of a false positive (at the specified *α* for the single-subject tests). The probability to pick a subject *i* from the population that shows an effect, assuming a population prevalence of *γ*, is thus

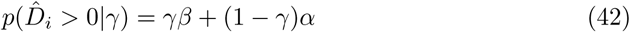

The probability to pick a subject from the population that shows no effect is

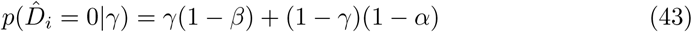

Thus the probability to observe *K* out of *N* subjects with an effect (see top panel) is

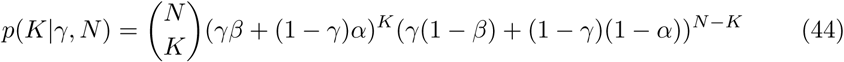

The sensitivity *β* is usually not known, and therefore is fixed at 1, to remain conservative. Computing the p-value as in Eq 41 to test the prevalence null hypothesis requires to sum over these values for *K* and higher and then to maximize over the range of *γ* values covered by the null hypothesis (see middle panel)

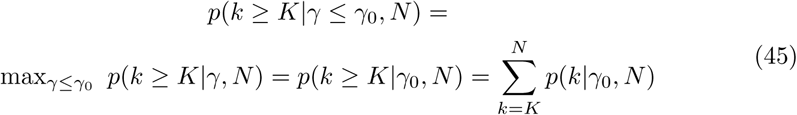

As discussed in (10), one can also report the largest *γ*_0_ value under which the null hypothesis can be rejected at the given significance level. This can be interpreted as the lower bound of a one-sided confidence interval about the true population prevalence *γ*, which can be of interest to the research question.

##### Statistical thresholding of subcorr maps across subjects

Statistical thresholding of the average *subcorr* maps computed from each subject’s signal subspace is obtained using permutation statistics, with the null hypothesis of exchangeability of signal and noise subspaces with respect to the statistic of interest. This latter is the average *subcorr* value across the group at a source location *ρ*.

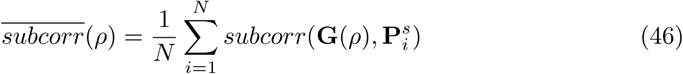

where**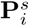** is the signal subspace of subject *i* in a group of *i* = 1, *…, N* subjects. If signal and noise subspaces are exchangeable with respect to the statistic of interest (the null hypothesis), we can randomly substitute the signal subspace with a dimensionality-matched noise subspace **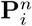** by selecting the columns of **P**_*i*_ associated with the smallest *D* values of the quality function *f*. We now run *O* permutations, where at each iteration we draw a binary permutation vector *ω* of length *N*. Then we compute the average *subcorr* value based on shuffled subspaces, where for each subject we use

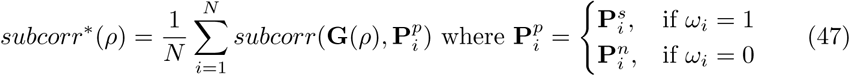

and keep the maximum *subcorr*^*\**^(*ρ*) over the volume at each iteration to obtain a null distribution against which to test the observed *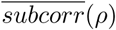*values across the volume.

## Supporting information

**S1 Fig.Additional contrasts and unthresholded maps for figure 5.** All analysis parameters are equivalent as described in the main part of the paper.

## Acknowledgments

PWD holds a Jeanne-Timmins Costello Fellowship from the Montreal Neurological Institute. S.B. is supported by a Discovery Grant from the National Science and Engineering Research Council of Canada (436355-13, http://www.nserc-crsng.gc.ca), the National Institutes of Health (2R01EB009048-05, http://nih.gov). The funders had no role in study design, data collection and analysis, decision to publish, or preparation of the manuscript.

## Footnotes

1 The code accompanying this paper can be found at https://github.com/pwdonh/ies_toolbox.

